# *uORF-Tools* – Workflow for the determination of translation-regulatory upstream open reading frames

**DOI:** 10.1101/415018

**Authors:** Anica Scholz, Florian Eggenhofer, Rick Gelhausen, Björn Grüning, Kathi Zarnack, Bernhard Brüne, Rolf Backofen, Tobias Schmid

## Abstract

Ribosome profiling (ribo-seq) provides a means to analyze active translation by determining ribosome occupancy in a transcriptome-wide manner. The vast majority of ribosome protected fragments (RPFs) resides within the protein-coding sequence of mRNAs. However, commonly reads are also found within the transcript leader sequence (TLS) (aka 5’ untranslated region) preceding the main open reading frame (ORF), indicating the translation of regulatory upstream ORFs (uORFs). Here, we present a workflow for the identification of translation-regulatory uORFs. Specifically, *uORF-Tools* identifies uORFs within a given dataset and generates a uORF annotation file. In addition, a comprehensive human uORF annotation file, based on 35 ribo-seq files, is provided, which can serve as an alternative input file for the workflow. To assess the translation-regulatory activity of the uORFs, stimulus-induced changes in the ratio of the RPFs residing in the main ORFs relative to those found in the associated uORFs are determined. The resulting output file allows for the easy identification of candidate uORFs, which have translation-inhibitory effects on their associated main ORFs. *uORF-Tools* is available as a free and open *Snakemake* workflow at https://github.com/Biochemistry1-FFM/uORF-Tools. It is easily installed and all necessary tools are provided in a version-controlled manner, which also ensures lasting usability. *uORF-Tools* is designed for intuitive use and requires only limited computing times and resources.

## Introduction

Translation is a highly regulated cellular process, regulation occurring predominantly at the level of initiation (Hinnebusch and Lorsch, 2012). Global translation is initiated in a cap-dependent manner, i.e. via binding of the cap-binding protein eukaryotic initiation factor 4E (eIF4E) to the 5’ 7-methyl-guanosine (m^7^G) cap present in all eukaryotic mRNAs and subsequent recruitment of the eIF4F initiation complex. As cap-dependent translation initiation depends on the availability of eIF4E, sequestration of the latter by eIF4E-binding proteins (4E-BPs), which are regulated by the central mTOR kinase, provides a means to efficiently control global translation (Thoreen *et al.*, 2012). In addition, there are numerous regulatory mechanisms that affect the translation of selected mRNAs only. Alternative modes of translational regulation commonly depend on *cis*-regulatory features within the 5’ untranslated region (UTR) of the respective mRNAs, e.g. specific sequences or secondary structures (Hinnebusch *et al.*, 2016). Alternative modes of translational regulation, such as internal ribosome entry site (IRES)-and upstream open reading frame (uORF)-dependent initiation, are of major importance under stress conditions, when global translation is inhibited, yet the synthesis of certain proteins needs to be sustained (Lacerda *et al.*, 2017; Walters and Thompson, 2016).

The analysis of translational changes was revolutionized by the development of the ribosome profiling (ribo-seq) technology, where actively translated regions are determined across the entire transcriptome by selective sequencing of ribosome protected footprints (RPFs) (Ingolia *et al.*, 2009). Sequencing reads in ribo-seq analyses are predominantly mapped to the protein-coding regions. Yet, while the 3’UTRs of transcripts usually lack RPFs, they are commonly observed in the 5’UTRs. Such actively translated regions are indicators for the presence of upstream open reading frames (uORFs), which represent short, peptide-coding sequences characterized by a start codon with an in-frame stop codon. Consequently, 5’UTRs are also referred to as transcript leader sequences (TLS) (Wethmar, 2014). With respect to their function, uORFs have been shown to affect the translation of associated main ORFs. While there are cases in which translation of a uORF positively affects the translation of the main ORF, for the most part efficient translation of a uORF is considered to restrict the translation of the respective main ORF (Young *et al.*, 2016). Of note, uORFs have been shown to play a prominent role during the integrated stress response (ISR), an adaptive response to various stress conditions aiming at restoring cellular homeostasis. During the ISR the translation initiation factor eIF2α is phosphorylated by protein kinase R (PKR), PKR-like endoplasmic reticulum stress (PERK), heme-regulated inhibitor (HRI), or general control non-repressible 2 (GCN2) kinases in response to stress conditions such as amino acid deprivation, viral infection, heme deprivation, and endoplasmic reticulum stress (Taniuchi *et al.*, 2016). Phosphorylation of eIF2α reduces global translation and at the same time enhances translation of selected, uORF-bearing mRNAs to allow for adaptation (Pakos-Zebrucka *et al.*, 2016). Such stress adaptive mechanisms are of major importance in a number of disease states including cancer and inflammation (Somers *et al.*, 2013).

There are various strategies to determine the presence of uORFs either based on sequence features within the TLS (McGillivray *et al.*, 2018), or using experimental ribo-seq data to identify actively translated ORFs including uORFs (Calviello *et al.*, 2015; Zhang *et al.*, 2017). With the present workflow, we aim to provide a pipeline that allows for the identification of differentially translated uORFs, which regulate the translation of the associated main ORFs. Using ribosome profiling data, *uORF-Tools* determines the experiment-specific, differentially translated uORFs and compares their translation with the translation of the respective main ORFs. While a uORF annotation file is generated for each individual experiment, a comprehensive human uORF annotation file, based on 35 data sets from nine human ribosome profiling data series, is also provided to allow for a comprehensive assessment of the translation regulatory impact of uORFs.

## Implementation and Workflow

### Implementation

*uORF-Tools* is provided as a free and open workflow and can be downloaded from https://github.com/Biochemistry1-FFM/uORF-Tools. It is based on *Snakemake* (Köster and Rahmann, 2012) and automatically installs all tool dependencies in a version-controlled manner via *bioconda* (Grüning *et al.*, 2018). The workflow can be run locally or in a cluster environment.

### Workflow

*uORF-Tools* is designed to receive bam files of ribosome profiling data sets as input (Figure 1). In addition, the workflow requires a genome fasta file and an annotation gtf file.

**Fig. 1.**
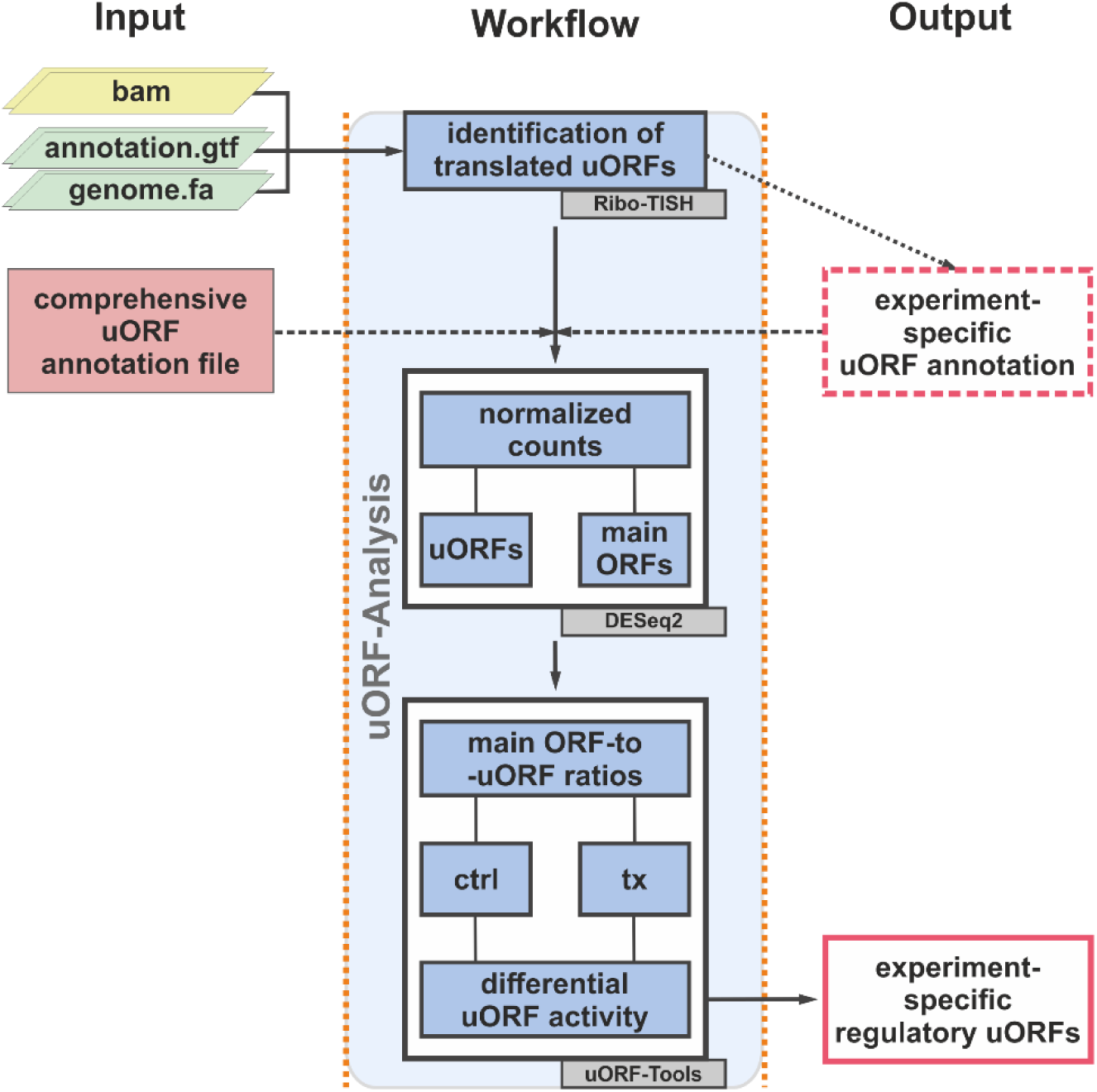
*uORF-Tools* – Workflow for the determination of translation-regulatory uORFs. Required input is shown on the left, a simplified depiction of processing in the center, and results on the right. (ctrl: control; tx: treatment)

Initially, *uORF-Tools* generates a new genome annotation file, which is used in the subsequent steps of the workflow. For practical reasons, this annotation file contains only the validated or manually annotated (confidence levels 1 and 2 in Gencode) (www.gencodegenes.org) longest protein coding transcript variants. Based on the provided input bam files and the generated genome annotation file, an experiment-specific uORF annotation file is then generated using *Ribo-TISH* (Zhang *et al.*, 2017). Specifically, *Ribo-TISH* identifies translation initiation sites within ribo-seq data and uses this information to determine ORFs, i.e. regular ORFs as well as uORFs. Default settings in uORF-Tools use the canonical start codon ATG only, yet users can allow for the use of alternative start codons as well. Furthermore, as uORFs are generally considered to be short, peptide-coding ORFs, a maximal length of 400 bp was set as upper size limit within the uORF-Tools pipeline for the identification of uORFs. To allow for an even broader characterization of potentially active uORFs, a comprehensive human uORF annotation file, based on 35 ribo-seq data sets, is provided with the package (for details see Supplementary Table S1). To use this comprehensive instead of the experiment-specific annotation file, the former needs to be selected by including its file path (uORF-Tools/comprehensive_annotation/uORF_annotation_hg38.csv) in the config.yaml file before starting the *uORF-Tools* workflow. Using uORF and genome annotation files, *uORF-Tools* creates one count file containing all reads that correspond to coding sequences (CDS) of the longest protein coding transcripts, i.e. main ORFs, and another count file which contains only reads that correspond to uORFs. To control for differences in library sizes, the count data are subsequently normalized using size factors calculated for all input libraries with *DESeq2* (Love *et al.*, 2014). To determine the relative translation of a main ORF, counts of the main ORF are normalized to the corresponding uORF counts. In order to assess if the main ORF-to-uORF ratios are altered in response to a stimulus, the differential uORF activity is determined by comparing the main ORF-to-uORF ratios between different conditions. A stimulus-dependent increase in the ratios indicates enhanced translation of the main ORF, i.e. reduced repression by the respective uORF, conversely a decrease in the ratios indicates that an inhibitory uORF becomes more active.

## Results and Discussion

*uORF-Tools* is provided as a readily deployable *Snakemake* workflow, which comes with extensive documentation (Supplementary Methods). Running *uORF-Tools* on 8 test data sets, i.e. 4 replicates of control and thapsigargin-treated HEK293 cells, with about 0.36 to 4.7 million reads per file on a consumer grade laptop (Intel® Core™ i5-8265U, 256 GB NVMe-SSD, 16 GB RAM) running Ubuntu 18.04.2 LTS required as little as 1.5 hours for a complete analysis. The input data and the utilized tools are clearly defined and enable reproducible analyses (Fig. 1; Supplementary Methods). Using an annotation.gtf and a genome.fa file obtained from Gencode (gencode.v28.annotation.gtf and GRCh38.p12.genome.fa), the analysis of the provided 8 test data sets (available at: ftp://biftp.informatik.uni-freiburg.de/pub/uORF-Tools/bam.tar.gzbam.tar.gz) identified 939 uORFs. In contrast, the provided, comprehensive uORF annotation file contains 1933 uORFs (Table 1, Supplementary Table S2). Interestingly, when the comprehensive annotation file was used in the analysis of the test data set, only 55 of the additional uORFs did not contain any RPF counts (Supplementary Table S3).

**Table 1.** Comparison of the performance of *uORF-Tools* for the 8 test data sets (GSE103719) using either the experiment-specific or the comprehensive annotation files.

|  | experiment-specific annotation file <sup>a</sup> | comprehensive annotation file |
| --- | --- | --- |
| identified uORFs | 939 | 1933 |
| mean length – main ORFs | 1509 | 1575 |
| mean length – uORFs | 38 | 40 |
| translation-inhibitory uORFs in test data set <sup>b</sup> | 47 | 94 |

(<sup>a</sup> using 8 test data sets (GSE103719); <sup>b</sup> 5% quantile of strongest changes in main ORF-to-uORF ratios)

To assess how the translation of the main ORFs might be affected by the uORFs, main ORF read counts were initially normalized to those of the associated uORFs (Figure 2A), which yielded mean ratios of 22.22 ± 4.56 or 28.69 ± 4.38 for the control and 27.19 ± 11.50 or 29.99 ± 10.89 for the thapsigargin-treated samples, based on the experiment-specific or the comprehensive annotations files, respectively.

**Fig. 2.**
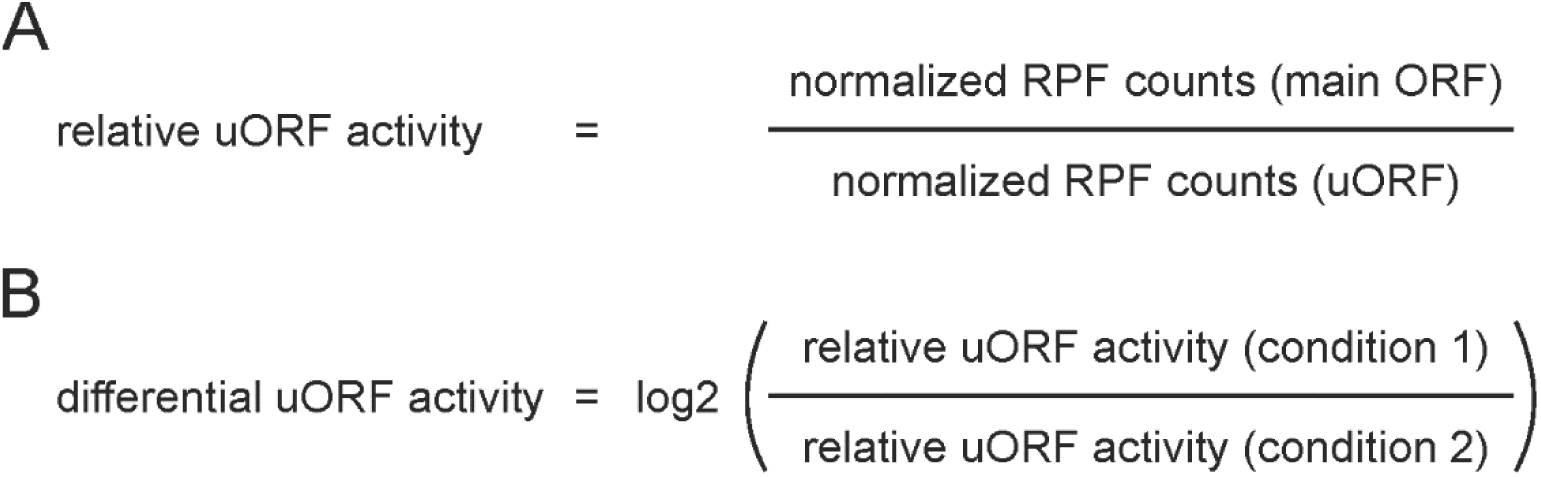
Calculation of uORF activities. (A) Relative uORF activities are determined for each experimental condition as ratio of the normalized ribosome protected fragment (RPF) counts of a specific main ORF relative the normalized RPF counts of the respective uORF. (B) Stimulus-dependent, differential uORF activities are then calculated as the log2 fold change of the ratio of the relative uORF activities of treatment (condition 1) vs. control (condition 2).

Despite the fact that the main ORFs of the uORF-bearing transcripts were generally much longer (mean lengths 1509 and 1575 bps, based on experiment-specific and comprehensive annotation files, respectively) than the uORFs (mean length 38 and 40 bps, based on experiment-specific and comprehensive annotation files, respectively) (Table 1), main ORF-to-uORF ratios as low as 0.018 were observed, indicating that some uORFs might be extremely potent in restricting the translation of their associated main ORF.

Subsequent calculation of the stimulus-dependent changes in main ORF-to-uORF ratios (Figure 2B), provides a means to easily identify uORFs inversely correlating with their associated main ORFs with respect to the transcript-specific ribosome occupancy.

In the case of the analyzed test data sets, the 5% quantile of the strongest changes in differential uORF activities was comprised of 47 transcripts based on the experiment-specific uORF annotation file, as compared to 94 transcripts in the case of the comprehensive uORF annotation file (Table 1, Supplementary Table S3). These differences underscore that it is advantageous to use comprehensive uORF annotations rather than experiment-specific ones only, as this might e.g. overcome low numbers of annotated uORFs due to ribo-seq analyses of either poor quality or containing low read numbers.

Along these lines, the experiment-specific annotation file identified only one uORF (uORF 2: 163-243 bp, 26 amino acids) within the TLS of the classical ISR target protein phosphatase 1 regulatory subunit 15A (PPP1R15A, aka growth arrest and DNA-damage-inducible 34 (GADD34)), whereas both published uORFs (uORF 1: 64-132 bp, 22 amino acids and uORF 2: 163-243 bp, 26 amino acids) (Lee *et al.*, 2009) were found using the comprehensive uORF annotation file. The uORFs within the TLS of PPP1R15A were exactly annotated as previously published. They were further found to be highly translated, as becomes already apparent when looking at the relative RPF peak heights of the uORFs relative to the main ORF, wherein a shift from a largely uORF-biased distribution under control conditions to more RPF reads in the thapsigargin group can be seen (Figure 3).

**Fig. 3.**
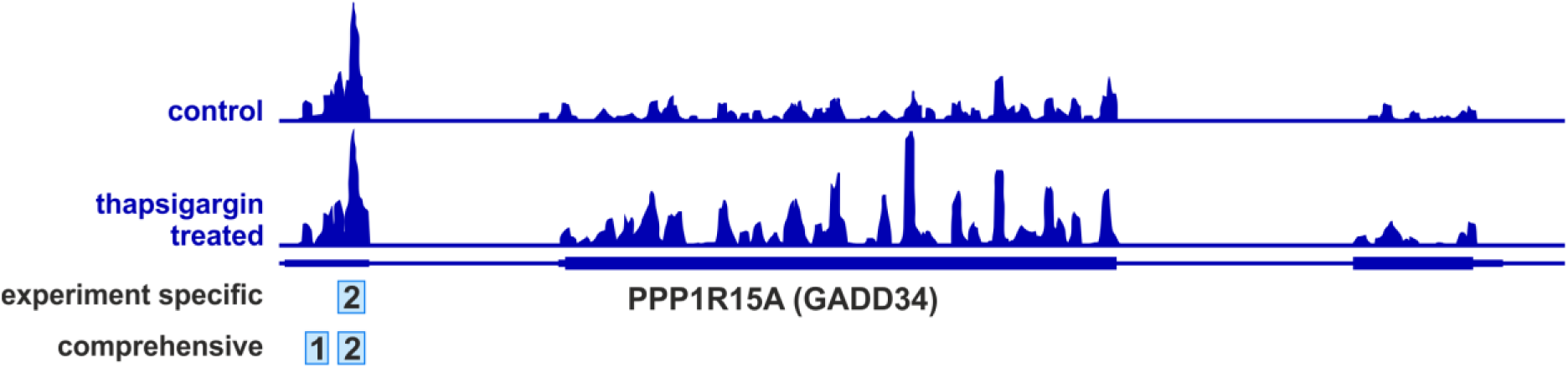
Distribution of RPF reads on the PPP1R15A (GADD34) transcript. Reads of control (*upper panel*) or thapsigargin-treated (*lower panel*) HEK293 from data set GSE103719 are shown. uORFs annotated either in the experiment-specific (2) or the comprehensive annotation file (1 and 2) within *uORF-Tools* are marked.

Quantitative analyses of the main ORF-to-uORF ratios revealed that the 5% quantile of strongest changes showed differential uORF activities of log2FC > |1.99| / |1.88| (based on the experiment-specific and the comprehensive annotation file, respectively) (Supplementary Tables S3).

In the case of PPP1R15A, translation under control conditions (main ORF-to-uORF ratios: uORF 1 = 7.85; uORF 2 = 4.54), shifted towards the main ORF under thapsigargin treatment (main ORF-to-uORF ratios: uORF 1 = 29.24; uORF 2 = 10.19). Not surprisingly, PPP1R15A emerged as one of the top candidates for uORF-dependent translational regulation in response to thapsigargin (Table 2). Specifically, the main ORF-to-uORF ratio of PPP1R15A displayed a log2FC increase of 1.87 and 4.26 for uORF 1 and uORF 2, respectively. This indicates that the translational repression under control conditions is relieved during the integrated stress response (ISR) and consequently the translation of the PPP1R15A main ORF increases, as previously reported (Lee *et al.*, 2009).

**Table 2.** *uORF-Tools* output. Top 10 candidates with the strongest increase or decrease in their main ORF-to-uORF ratios (mean reads > 10 for each condition and ORF as well as uORF).

| Coordinates | Gene symbol | Transcript id | uORF id | mean reads<br>uORF<br>(con. 1) | mean reads<br>ORF<br>(con. 1) | mean reads<br>uORF<br>(con. 2) | mean reads<br>ORF<br>(con. 2) | Log2FC<br>(main ORF-to-uORF ratios) |
| --- | --- | --- | --- | --- | --- | --- | --- | --- |
| chr5:<br>112976904-<br>112985968:+ | DCP2 | ENST00000<br>389063.2 | ENST00000<br>389063.2.1 | 25.94 | 92.42 | 49.70 | 98.60 | 3.39 |
| chr7:<br>87858731-<br>87876195:- | SLC25A40 | ENST00000<br>341119.9 | ENST00000<br>341119.9.1 | 13.10 | 30.94 | 10.37 | 20.83 | 2.56 |
| chr1:<br>154272679-<br>154272727:+ | HAX1 | ENST00000<br>328703.11 | ENST00000<br>328703.11.1 | 12.69 | 114.73 | 20.68 | 98.93 | 2.54 |
| chr17:<br>77089165-<br>77093272:+ | SEC14L1 | ENST00000<br>392476.6 | ENST00000<br>392476.6.2 | 104.09 | 55.69 | 22.75 | 85.23 | -2.53 |
| chr5:<br>128537690-<br>128539509:- | FBN2 | ENST00000<br>508053.5 | ENST00000<br>508053.5.1 | 16.48 | 460.77 | 15.93 | 494.25 | 2.40 |
| chr19:<br>17555646-<br>17559337:+ | GLT25D1 | ENST00000<br>252599.8 | ENST00000<br>252599.8.1 | 11.83 | 93.63 | 20.40 | 113.72 | 2.38 |
| chr7:<br>158693469-<br>158701925:- | NCAPG2 | ENST00000<br>356309.7 | ENST00000<br>356309.7.1 | 10.24 | 175.92 | 13.95 | 196.84 | 2.26 |
| chr19:<br>48872553-<br>48872634:+ | PPP1R15A | ENST00000<br>200453.5 | ENST00000<br>200453.5.1 | 40.37 | 411.77 | 10.55 | 47.95 | 1.87 |
| chr12:<br>47964211-<br>47964298:+ | TMEM106<br>C | ENST00000<br>256686.10 | ENST00000<br>256686.10.1 | 12.56 | 55.26 | 18.97 | 68.44 | 1.80 |
| chr7:<br>35801209-<br>35832814:+ | SEPT7 | ENST00000<br>399034.7 | ENST00000<br>399034.7.1 | 15.91 | 62.62 | 18.60 | 88.40 | 1.76 |

Corroborating the concept that the translation of otherwise uORF-repressed main ORFs is elevated during the ISR, only 6 of the 94 candidates within the 5% quantile of strongest changes, as identified using the comprehensive annotation file, showed reduced main ORF-to-uORF ratios, i.e. enhanced translational repression by the uORF. Furthermore, 675 candidates had main ORF-to-uORF ratios log2FC < 0 (73 candidates log2FC < −1), while 1245 had log2FC > 0 (411 candidates log2FC > 1). Along the same lines, the mean of all reduced main ORF-to-uORF ratios was log2FC = −0.50 and the mean for all elevated ones log2FC = 0.83 (Supplementary Table S3). All of these findings support the notion that thapsigargin relieves uORF-mediated translational repression of specific targets.

In addition to the identification of translation-inhibitory uORFs, the output file also contains uORFs that are regulated in the same direction as their associated main ORFs, which may indicate a translation-supportive function of the respective uORFs. The candidates within the 5% quantile of least changes display main ORF-to-uORF ratios log2FC < |0.05|. It should be noted that, while the translation-inhibitory uORFs are easily identified with *uORF-Tools*, the unambiguous identification of translation-supporting uORFs would require additional information such as the translation efficiencies.

In addition to the stimulus-dependent changes in main ORF-to-uORF ratios, as indicators for translation regulatory uORF activities, the final output table provides the mean read counts for uORFs and main ORFs under all conditions tested (Table 2, Supplementary Table S3). This will be informative for the assessment of the translational status of the individual transcripts and, thus, the potential relevance of the determined changes in a given data set.

## Availability and Future Directions

The *uORF-Tools* workflow is provided as free and open software (https://github.com/Biochemistry1-FFM/uORF-Tools), which can be easily deployed with all version-controlled dependencies. Bam files of the used 8 test data sets (GSE103719; Woo *et al.*, 2018) are available at ftp://biftp.informatik.uni-freiburg.de/pub/uORF-Tools/bam.tar.gz. Extensive documentation of the workflow is provided with the software and supplied in the Supplementary Methods.

*uORF-Tools* generates an intuitive, easy to interpret output file, containing the stimulus-dependent changes in main ORF-to-uORF ratios as an indicator for uORFs that negatively regulate the translation of their associated main ORFs. In addition, *uORF-Tools* provides a comprehensive human uORF annotation file based on 35 ribosome profiling data sets (Supplementary Table 1), which appeared superior to experiment-specific uORF annotation files, with respect to the identification of translation-regulatory uORFs.

Even with limited computing resources, *uORF-Tools* is a fast software solution and a valuable addition to the portfolio of methods for researchers interested in the function of uORFs.

## Supporting Information Legends

### Supplementary Methods

Supplementary Methods gives a more detailed description of the *uORF-Tools* workflow and its implementation.

**Supplementary Table S1.** Ribo-seq data series used for the generation of the comprehensive human uORF annotation file. 35 data sets from nine different data series each included ribo-seq and associated RNA-seq data.

**Supplementary Table S2.** Lists of uORFs in the comprehensive and in the experiment-specific annotation files.

**Supplementary Table S3.** Output files generated using the comprehensive or the experiment-specific annotation files.

## References

Calviello L, Mukherjee N, Wyler E, Zauber H, Hirsekorn A, Selbach M, et al. Detecting actively translated open reading frames in ribosome profiling data. Nat Methods. 2016;13(2): 165–170.

Grüning B, Dale R, Sjödin A, Chapman BA, Rowe J, Tomkins-Tinch CH, et al. Bioconda: sustainable and comprehensive software distribution for the life sciences. Nat Methods. 2018;15(7): 475–476.

Hinnebusch AG, Lorsch JR. The mechanism of eukaryotic translation initiation: new insights and challenges. Cold Spring Harb Perspect Biol. 2012;4(10): a011544.

Hinnebusch AG, Ivanov IP, Sonenberg N. Translational control by 5′-untranslated regions of eukaryotic mRNAs. Science. 2016;352(6292): 1413–1416.

Ingolia NT, Ghaemmaghami S, Newman JR, Weissman JS. Genome-wide analysis *in vivo* of translation with nucleotide resolution using ribosome profiling. Science. 2009;324(5924): 218–223.

Köster J, Rahmann S. Snakemake--a scalable bioinformatics workflow engine. Bioinformatics. 2012;28(19): 2520–2522.

Lacerda R, Menezes J, Romão L. More than just scanning: the importance of cap-independent mRNA translation initiation for cellular stress response and cancer. Cell Mol Life Sci. 2017;74(9): 1659–1680.

Lee Y, Cevallos RC, Jan E. An Upstream Open Reading Frame Regulates Translation of GADD34 during Cellular Stresses That Induce eIF2α Phosphorylation. J Biol Chem. 2009; 284: 6661–6673.

Love MI, Huber W, Anders S. Moderated estimation of fold change and dispersion for RNA-seq data with DESeq2. Genome Biol. 2014;15(12): 550.

McGillivray P, Ault R, Pawashe M, Kitchen R, Balasubramanian S, Gerstein M. A comprehensive catalog of predicted functional upstream open reading frames in humans. Nucleic Acids Res. 2018;46(7): 3326–3338.

Pakos-Zebrucka K, Koryga I, Mnich K, Ljujic M, Samali A, Gorman AM. The integrated stress response. EMBO Rep. 2016;17(10): 1374–1395.

Somers J, Pöyry T, Willis AE. A perspective on mammalian upstream open reading frame function. Int J Biochem Cell Biol. 2013;45(8): 1690–1700.

Taniuchi S, Miyake M, Tsugawa K, Oyadomari M, Oyadomari S. Integrated stress response of vertebrates is regulated by four eIF2α kinases. Sci Rep. 2016;6:32886.

Thoreen CC, Chantranupong L, Keys HR, Wang T, Gray NS, Sabatini DM. A unifying model for mTORC1-mediated regulation of mRNA translation. Nature. 2012;485(7396): 109–113.

Walters B, Thompson SR. Cap-Independent Translational Control of Carcinogenesis. Front Oncol. 2016;6:128.

Wethmar K. The regulatory potential of upstream open reading frames in eukaryotic gene expression. Wiley Interdiscip Rev RNA. 2014;5(6): 765–778.

Woo YM, Kwak Y, Namkoong S, Kristjánsdóttir K, Lee SH, Lee JH, et al. TED-Seq Identifies the Dynamics of Poly(A) Length during ER Stress. Cell Rep. 2018;24(13): 3630–3641.

Young SK, Wek RC. Upstream Open Reading Frames Differentially Regulate Gene-specific Translation in the Integrated Stress Response. J Biol Chem. 2016;291(33): 16927–16935.

Zhang P, He D, Xu Y, Hou J, Pan BF, Wang Y, et al. Genome-wide identification and differential analysis of translational initiation. Nat Commun. 2017;8(1): 1749.

