## Supplementary Methods for "*uORF-Tools* – Workflow for the determination of translation-regulatory upstream open reading frames"

### Workflow documentation

### 1 Introduction

uORF-Tools is a workflow and a collection of tools for the analysis of 'Upstream Open Reading Frames' (short uORFs). The workflow is based on the workflow management system *snakemake* [1] and handles installation of all dependencies via *bioconda* [2], as well as all processings steps. The source code of *uORF-Tools* is open source and available under the License *GNU General Public License 3*. Installation and basic usage is described below.
**This documentation was last updated 08.03.2019. An up-to-date web-based version of the documentation can be found on ReadTheDocs.**

### 2 Program flowchart

The flowchart (see Figure 1) describes the processing steps of the workflow and how they are connected.

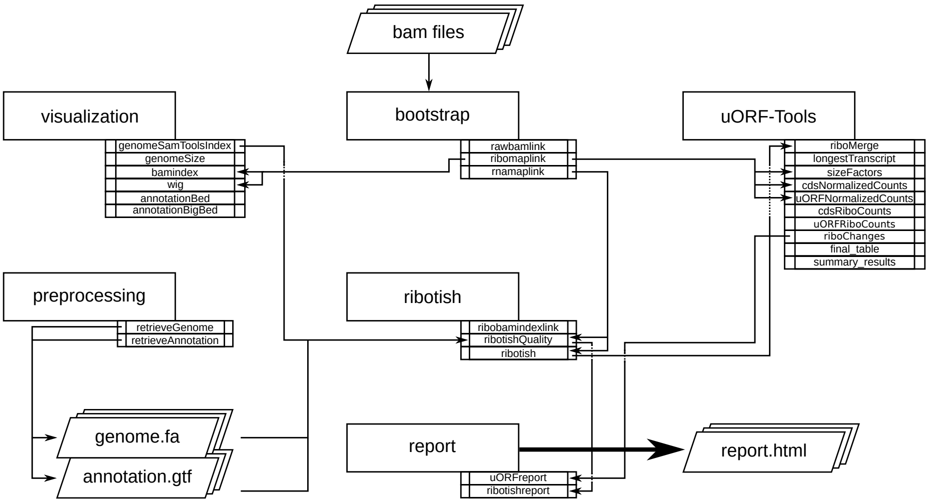

**Fig. 1.** Sketch of the workflow.

#### 3 Directory table

The output is written to a directory structure (see Figure 2) that corresponds to the workflow steps.

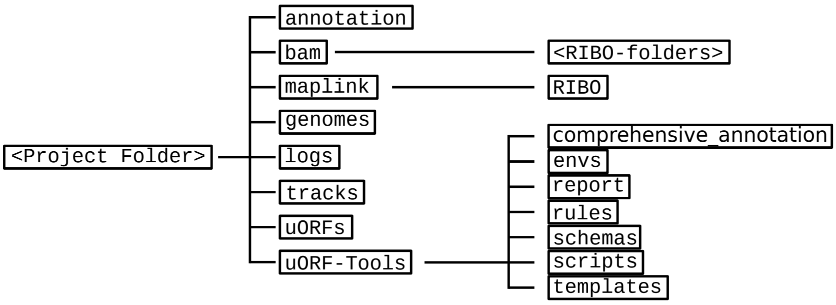

**Fig. 2.** The directory structure created by running the workflow.

- **annotation**: contains the processed user-provided annotation file with genomic features.
  Contents: *annotation.gtf*
- **bam**: contains the input *.bam* files.
  Contents: *<method-condition-replicate>.bam*
- **genomes**: contains the genome file, as well as an according index and sizes file.
  Contents: *genome.fa*, *genome.fa.fai*, *sizes.genome*
- **index**: contains an index for the rRNA databases created using indexdb_rna.
  Contents: *<database-ID>.bursttrie_0.dat*, *<database-ID>.kmer_0.dat*, *<database-ID>.pos_0.dat*, *<database-ID>.stats*
- **logs**: contains log files for each step of the workflow.
  Contents: *<rule>.o<jobID>*, *<methods>.log*
- **maplink**: contains soft links to the *.bam* files and an according index.
  - **RIBO**: contains soft links to the *.bam* and *.bam.bai* files for RIBO and corresponding parameter files (*.para.py*).

  Contents: *<method-condition-replicate>.bam.bai*, *RIBO/<condition-replicate>.bam.para.py*
- **tracks**: contains *BED (.bed)*, *wig (.wig)* and *bigWig (.bw)* files for visualizing tracks in a genome browser.
  Contents: *annotation.bb*, *annotation.bed*, *annotation.bed6*, *annotationNScore.bed6*, *annotation-woGenes.gtf*, *<method-condition-replicate>.bw* *<method-condition-replicate>.wig*

- **uORFs**: contains the main output of the workflow.
  - **uORFs_regulation.tsv**: table summarizing the predicted uORFs with their regulation on the main ORF.
  - **merged_uORFs.bed**: genome browser track with predicted uORFs.

  Contents: *longest_protein_coding_transcripts.gtf*, *uORF_regulation.tsv*, *ribo_norm_CDS_reads.csv*, *ribo_norm_uORFs_reads.csv*, *ribo_raw_CDS_reads.csv*, *ribo_raw_uORFs_reads.csv*, *sfactors_lprot.csv*, *merged_uORFs.bed*, *merged_uORFs.csv*,
- **uORF-Tools**: contains the workflow tools.
  - **comprehensive_annotation**: an example annotation.
  - **envs**: conda environment files (.yaml).
  - **report**: restructuredText files for the report (.rst).
  - **rules**: the snakemake rules.
  - **schemas**: validation templates for input files
  - **scripts**: scripts used by the snakemake workflow.
  - **templates**: templates for the *config.yaml* and the *samples.tsv*.

### 4 Requirements

In the following, we describe all the required files and tools needed to run our workflow.

#### 4.1 Tools

##### miniconda3

As this workflow is based on the workflow management system *snakemake* [1]. *Snakemake* will download and install all necessary dependencies via *conda*.
We strongly recommend installing *miniconda3* with *python3.7*.
After downloading the *miniconda3* version suiting your linux system, execute the downloaded bash file and follow the instructions given.

##### snakemake

**The uORF-Tools require *snakemake* (version == 5.4.2).**
The newest version of *snakemake* can be downloaded via conda using the following command:

```
$ conda create -c conda-forge -c bioconda \
               -n snakemake snakemake==5.4.2
```

This creates a new conda environment called *snakemake* and installs *snakemake* into the environment.
**Due to conflicts between newer conda and snakemake versions, the workflow relies on an older version of conda. Simply update your conda using:**.

```
$ conda install -n base conda=4.5.13
```

The environment can be activated using:

```
$ source activate snakemake
```

and deactivated using:

```
$ source deactivate
```

##### uORF-Tools

Using the workflow requires the *uORF-Tools*. The latest version is available on our GitHub page.
In order to run the workflow, we suggest that you download the *uORF-Tools* into your project directory. The following command creates an example directory and changes into it:

```
$ mkdir project; cd project;
```

Now, download and unpack the latest version of the *uORF-Tools* by entering the following commands:

```
$ wget https://github.com/Biochemistry1-FFM\
/uORF-Tools/archive/3.0.0.tar.gz
$ tar -xzf 3.0.0.tar.gz; mv uORF-Tools-3.0.0 uORF-Tools;
$ rm 3.0.0.tar.gz;
```

The *uORF-Tools* are now located in a subdirectory of your workflow directory.

#### 4.2 Input Files

Several input files are required in order to run our workflow, a genome sequence (.fa), an annotation file (.gtf) and the bam files (.bam).

##### genome.fa and annotation.gtf

We recommend retrieving both the genome and the annotation files for mouse and human from GENCODE [3] and for other species from Ensembl Genomes [4].

##### input .bam files

These are the input files provided by you (the user).
"For best performance, reads should be trimmed (to ~ 29 nt RPF length) and aligned to genome using end-to-end mode (no soft-clip). Intron splicing is supported. Some attributes are needed such as NM, NH and MD. For STAR, '–outSAMattributes All' should be set. bam file should be sorted and indexed by samtools." (RiboTISH requirements, see https://github.com/zhpn1024/ribotish' ).
Please ensure that you move all input *.bam* files into a folder called *bam*:

```
$ mkdir bam
$ mv *.bam bam/
```

**sample sheet and configuration file**

In order to run the *uORF-Tools*, you have to provide a sample sheet and a configuration file. There are templates for both files available in the *uORF-Tools* folder.
Copy the templates of the sample sheet and the configuration file into the *uORF-Tools* folder:

```
$ cp uORF-Tools/templates/samples.tsv uORF-Tools/
$ cp uORF-Tools/templates/config.yaml uORF-Tools/
```

Customize the *config.yaml* using your preferred editor. It contains the following variables:

- **taxonomy** Specify the taxonomic group of the used organism in order to ensure the correct removal of reads mapping to ribosomal genes (Eukarya, Bacteria, Archea). (Option for the preprocessing workflow)
- **adapter** Specify the adapter sequence to be used. If not set, *Trim galore* will try to determine it automatically. (Option for the preprocessing workflow)
- **samples** The location of the samples sheet created in the previous step.
- **genomeindexpath** If the STAR genome index was already precomputed, you can specify the path to the files here, in order to avoid recomputation. (Option for the extended workflow)
- **uorfannotationpath** If a uORF-annotation file was already pre-computed, you can specify the path to the file here. Please make sure, that the file has the same format as the uORF_annotation_hg38.csv file provided in the git repo (i.e. same number of columns, same column names)
- **alternativestartcodons** Specify a comma separated list of alternative start codons.

Edit the sample sheet corresponding to your project. It contains the following variables:

- **method** Indicates the method used for this project, here RIBO for ribosome profiling.
- **condition** Indicates the applied condition (A, B)
- **replicate** Used to distinguish between the different replicates (e.g. 1,2, ...)
- **inputFile** Indicates the bam file for a given sample.

As seen in the *samples.tsv* template:
**Please make sure that you have at-least two replicates for each condition!**.
**Please ensure that you put the treatment before the control alphabetically (e.g. A: Treatment B: Control)**.

| method | condition | replicate | inputFile |
|---|---|---|---|
| RIBO | A | 1 | bam/RIBO-A-1.bam |
| RIBO | A | 2 | bam/RIBO-A-2.bam |
| RIBO | A | 3 | bam/RIBO-A-3.bam |
| RIBO | A | 4 | bam/RIBO-A-4.bam |
| RIBO | B | 1 | bam/RIBO-B-1.bam |
| RIBO | B | 2 | bam/RIBO-B-2.bam |
| RIBO | B | 3 | bam/RIBO-B-3.bam |
| RIBO | B | 4 | bam/RIBO-B-4.bam |

##### cluster.yaml

In the *template* folder, we provide two cluster.yaml files needed by snakemake in order to run on a cluster system:

- **sge-cluster.yaml** for grid based queuing systems
- **torque-cluster.yaml** for torque based queuing systems

### 5 Example workflow

The retrieval of input files and running the workflow locally and on a server cluster via a queuing system is demonstrated using an example with data available from our FTP-Server.
**Ensure that you have miniconda3 installed and a conda environment set-up. Please refer to the Tools Section (4.1) for details on the installation.**

#### 5.1 Setup

First of all, we start by creating the project directory and changing to it.

```
$ mkdir project; cd project;
```

We then download the latest version of the *uORF-Tools* into the newly created project folder and unpack it.

```
$ wget https://github.com/Biochemistry1-FFM\
/uORF-Tools/archive/3.0.0.tar.gz
$ tar -xzf 3.0.0.tar.gz; mv uORF-Tools-3.0.0 uORF-Tools;
$ rm 3.0.0.tar.gz;
```

#### 5.2 Retrieve and prepare input files

Before starting the workflow, we have to acquire and prepare several input files. These files are the annotation file, the genome file, the bam files, the configuration file and the sample sheet.

##### Annotation and genome files

First, we want to retrieve the annotation file and the genome file. In this case, we can find both on the GENCODE [3] webpage for the human genome.
On this page, we can directly retrieve both files by clicking on the according download links next to the file descriptions. (As shown in Figure 3). Alternatively, you can directly download them using the following commands:

```
$ wget ftp://ftp.ebi.ac.uk/pub/databases/gencode\
     /Gencode_human/release_28/gencode.v28.annotation.gtf.gz
$ wget ftp://ftp.ebi.ac.uk/pub/databases/gencode\
     /Gencode_human/release_28/GRCh38.p12.genome.fa.gz
```

GTF / GFF3 files

| Content | Regions | Description | Download |
|---|---|---|---|
| Comprehensive gene annotation | CHR | • It contains the comprehensive gene annotation on the reference chromosomes only<br>• This is the **main annotation file** for most users | GTF GFF3 |
| Comprehensive gene annotation | ALL | • It contains the comprehensive gene annotation on the reference chromosomes, scaffolds, assembly patches and alternate loci (haplotypes)<br>• This is a **superset** of the main annotation file | GTF GFF3 |

Fasta files

| Content | Regions | Description | Download |
|---|---|---|---|
| Transcript sequences | CHR | • Nucleotide sequences of all transcripts on the reference chromosomes | Fasta |
| Protein-coding transcript sequences | CHR | • Nucleotide sequences of coding transcripts on the reference chromosomes<br>• Transcript biotypes: protein_coding, nonsense_mediated_decay, non_stop_decay, IG_*_gene, TR_*_gene, polymorphic_pseudogene | Fasta |
| Protein-coding transcript translation sequences | CHR | • Amino acid sequences of coding transcript translations on the reference chromosomes<br>• Transcript biotypes: protein_coding, nonsense_mediated_decay, non_stop_decay, IG_*_gene, TR_*_gene, polymorphic_pseudogene | Fasta |
| Long non-coding RNA transcript sequences | CHR | • Nucleotide sequences of long non-coding RNA transcripts on the reference chromosomes | Fasta |
| Genome sequence (GRCh38.p12) | ALL | • Nucleotide sequence of the GRCh38.p12 genome assembly version on all regions, including reference chromosomes, scaffolds, assembly patches and haplotypes<br>• The sequence region names are the same as in the GTF/GFF3 files | Fasta |

**Fig. 3.** Downloading the annotation file and the genome file from the GenCode webpage using the direct download links.

Then, we are going to unpack both files.

```
$ gunzip gencode.v28.annotation.gtf.gz
$ gunzip GRCh38.p12.genome.fa.gz
```

Finally, we will rename these files to *annotation.gtf* and *genome.fa*.

```
$ mv gencode.v28.annotation.gtf annotation.gtf
$ mv GRCh38.p12.genome.fa genome.fa
```

Another webpage that provides these files is Ensembl Genomes. This usually requires searching their file system in order to find the wanted files. For this tutorial, we recommend to stick to GenCode instead.

##### .bam files

Next, we want to acquire the bam files. The bam files for the tutorial dataset can be downloaded from our FTP-Server:
**We provide both a .zip and a .tar.gz file. We recommend the .tar.gz file as most linux systems can decompress them via commandline by default.**

```
$ wget ftp://biftp.informatik.uni-freiburg.de/
     \pub/uORF-Tools/bam.tar.gz;
$ tar -zxvf bam.tar.gz; rm bam.tar.gz;
```

This will create a bam folder containing all the files necessary to run the workflow. If you prefer using your own .bam files, we suggest creating a bam folder and copying the files into it. Make sure that your reads were trimmed (to ~29bp length) and aligned to the genome using end-to-end alignment. The bam files need to include all SAM attributes and should be sorted using samtools.

```
$ mkdir bam; mv *.bam bam/;
```

##### Configuration file and sample sheet

Finally, we will prepare the configuration file (*config.yaml*) and the sample sheet (*samples.tsv*). We start by copying templates for both files from the *uORF-Tools/templates* into the *uORF-Tools/* folder.

```
$ cp uORF-Tools/templates/bam-samples.tsv uORF-Tools/
$ mv uORF-Tools/bam-samples.tsv uORF-Tools/samples.tsv
```

The standard template *bam-samples.tsv* looks as follows:

| method | condition | replicate | inputFile |
|---|---|---|---|
| RIBO | A | 1 | bam/RIBO-A-1.bam |
| RIBO | A | 2 | bam/RIBO-A-2.bam |
| RIBO | A | 3 | bam/RIBO-A-3.bam |
| RIBO | A | 4 | bam/RIBO-A-4.bam |
| RIBO | B | 1 | bam/RIBO-B-1.bam |
| RIBO | B | 2 | bam/RIBO-B-2.bam |
| RIBO | B | 3 | bam/RIBO-B-3.bam |
| RIBO | B | 4 | bam/RIBO-B-4.bam |

**When using your own data, use any editor (vi(m), gedit, nano, atom, ...) to customize the sample sheet.**

**Please ensure not to replace any tabulator symbols with spaces while changing this file.**

Next, we are going to set up the *config.yaml*.

```
$ cp uORF-Tools/templates/config.yaml uORF-Tools/
$ vi uORF-Tools/config.yaml
```

This file contains the following variables:

- **taxonomy** Specify the taxonomic group of the used organism in order to ensure the correct removal of reads mapping to ribosomal genes (Eukarya, Bacteria, Archea). (Option for the preprocessing workflow)
- **adapter** Specify the adapter sequence to be used. If not set, *Trim galore* will try to determine it automatically. (Option for the preprocessing workflow)
- **samples** The location of the samples sheet created in the previous step.
- **genomeindexpath** If the STAR genome index was already precomputed, you can specify the path to the files here, in order to avoid recomputation. (Option for the preprocessing workflow)

- **uorfannotationpath** If a uORF-annotation file was already pre-computed, you can specify the path to the file here. Please make sure, that the file has the same format as the uORF_annotation_hg38.csv file provided in the git repo (i.e. same number of columns, same column names)
- **alternativestartcodons** Specify a comma separated list of alternative start codons.

```
#Taxonomy of the samples to be processed, possible are Eukarya
    , Bacteria, Archea
taxonomy: "Eukarya"
#Adapter sequence used
adapter: ""
samples: "uORF-Tools/samples.tsv"
genomeindexpath: ""
uorfannotationpath: ""
alternativestartcodons: "CTG,GTG,TTG"
```

**Fig. 4.** The contents of the configuration file (*config.yaml*)

For this tutorial, we can keep the default values for the *config.yaml* as shown in Figure 6. The organism analyzed in this tutorial is *homo sapiens*, therefore we keep the taxonomy at *Eukarya*. The path to *samples.tsv* is set correctly.

#### 6 Running the workflow

Now that we have all the required files, we can start running the workflow, either locally or in a cluster environment.

**Run the workflow locally** Use the following steps when you plan to execute the workflow on a single server or workstation. Please be aware that some steps of the workflow require a lot of memory, specifically for eukaryotic species.

```
$ snakemake --use-conda -s uORF-Tools/Snakefile \
            --configfile uORF-Tools/config.yaml \
            --directory ${PWD} -j 20 \
            --latency-wait 60
```

**Run Snakemake in a cluster environment** Use the following steps if you are executing the workflow via a queuing system. Edit the configuration file cluster.yaml according to your queuing system setup and cluster hardware. The following system call shows the usage with Grid Engine:

```
$ snakemake --use-conda -s uORF-Tools/Snakefile \
       --configfile uORF-Tools/config.yaml \
       --directory ${PWD} -j 20 \
       --cluster-config uORF-Tools/templates/sge-cluster.yaml
```

```
#!/bin/bash
#PBS -N <ProjectName>
#PBS -S /bin/bash
#PBS -q "long"
#PBS -d <PATH/ProjectFolder>
#PBS -l nodes=1:ppn=1
#PBS -o <PATH/ProjectFolder>
#PBS -j oe
cd <PATH/ProjectFolder>
source activate snakemake
snakemake --latency-wait 600 --use-conda -s uORF-Tools/
    Snakefile --configfile uORF-Tools/config.yaml --
    directory ${PWD} -j 20 --cluster-config uORF-Tools/
    torque.yaml --cluster "qsub -N {cluster.jobname} -S /
    bin/bash -q {cluster.qname} -d <PATH/ProjectFolder> -l
     {cluster.resources} -o {cluster.logoutputdir} -j oe"
```

**Fig. 5.** Bash script to run the example workflow in a TORQUE cluster environment.

**Example: Run Snakemake in a cluster environment** We ran the tutorial workflow in a cluster environment, specifically a TORQUE cluster environment. Therefore, we created a bash script *torque.sh* in our project folder.

```
$ vi torque.sh
```

We proceeded by writing the queueing script as shown in Figure 7.
We then simply submitted this job to the cluster.

```
$ qsub torque.sh
```

Using any of the presented methods, this will run the workflow on our dataset and create the desired output files. Once the workflow has finished, we can request an automatically generated *report.html* file using the following command:

```
$ snakemake --latency-wait 600 --use-conda \
            -s uORF-Tools/Snakefile \
            --configfile uORF-Tools/config.yaml \
            --report report.html
```

#### 7 Preprocessing workflow

The retrieval of input files and running the workflow locally and on a server cluster via a queuing system is demonstrated using an example with data available from SRA via NCBI. The dataset is available under the GEO accession number GSE103719. The retrieval of the data is described in this tutorial.
**Ensure that you have miniconda3 installed and a conda environment set-up. Please refer to the Tools Section (4.1) for details on the installation.**

##### 7.1 Setup

First of all, we start by creating the project directory and changing to it.

```
$ mkdir preprocessing_project; cd preprocessing_project;
```

We then download the latest version of the *uORF-Tools* into the newly created project folder and unpack it.

```
$ wget https://github.com/Biochemistry1-FFM\
/uORF-Tools/archive/3.0.0.tar.gz
$ tar -xzf 3.0.0.tar.gz; mv uORF-Tools-3.0.0 uORF-Tools;
$ rm 3.0.0.tar.gz;
```

##### 7.2 Retrieve and prepare input files

Before starting the workflow, we have to acquire and prepare several input files. These files are the annotation file, the genome file, the fastq files, the configuration file and the sample sheet.

##### Annotation and genome files

On this page, we can directly retrieve both files by clicking on the according download links next to the file descriptions. (As shown in Figure 3). Alternatively, you can directly download them using the following commands:

```
$ wget ftp://ftp.ebi.ac.uk/pub/databases/gencode\
     /Gencode_human/release_28/gencode.v28.annotation.gtf.gz
$ wget ftp://ftp.ebi.ac.uk/pub/databases/gencode\
     /Gencode_human/release_28/GRCh38.p12.genome.fa.gz
```

Then, we are going to unpack both files.

```
$ gunzip gencode.v28.annotation.gtf.gz
$ gunzip GRCh38.p12.genome.fa.gz
```

Finally, we will rename these files to *annotation.gtf* and *genome.fa*.

```
$ mv gencode.v28.annotation.gtf annotation.gtf
$ mv GRCh38.p12.genome.fa genome.fa
```

Another webpage that provides these files is Ensembl Genomes. This usually requires searching their file system in order to find the wanted files. For this tutorial, we recommend to stick to GenCode instead.

##### Fastq files

In this example, we will use both RNA-seq and RIBO-seq data. In order to fasten up the tutorial, we download only 2 of the 4 replicates available for each Condition.
**Please note that you should always use all available replicates, when analyzing your data.**

##### European Nucleotide Archive (ENA)

For many datasets, the easiest way to retrieve the fastq files is using the European Nucleotide Archive as it provides direct download links when searching for a dataset. Use the interface on ENA or type the follwing commands:

```
$ wget ftp://ftp.sra.ebi.ac.uk/vol1/fastq/\
           SRR602/005/SRR6026765/SRR6026765.fastq.gz;
$ wget ftp://ftp.sra.ebi.ac.uk/vol1/fastq/\
           SRR602/005/SRR6026765/SRR6026766.fastq.gz;
$ wget ftp://ftp.sra.ebi.ac.uk/vol1/fastq/\
           SRR602/005/SRR6026765/SRR6026769.fastq.gz;
$ wget ftp://ftp.sra.ebi.ac.uk/vol1/fastq/\
           SRR602/005/SRR6026765/SRR6026770.fastq.gz;
$ wget ftp://ftp.sra.ebi.ac.uk/vol1/fastq/\
           SRR602/005/SRR6026765/SRR6026773.fastq.gz;
```

```
$ wget ftp://ftp.sra.ebi.ac.uk/vol1/fastq/\
           SRR602/005/SRR6026765/SRR6026774.fastq.gz;
$ wget ftp://ftp.sra.ebi.ac.uk/vol1/fastq/\
           SRR602/005/SRR6026765/SRR6026777.fastq.gz;
$ wget ftp://ftp.sra.ebi.ac.uk/vol1/fastq/\
           SRR602/005/SRR6026765/SRR6026778.fastq.gz;
```

Then, we create a fastq folder and move all the *.fastq.gz* files into this folder.

```
$ mkdir fastq; mv *.fastq.gz fastq/;
```

##### Configuration file and sample sheet

Finally, we will prepare the configuration file (*config.yaml*) and the sample sheet (*samples.tsv*). We start by copying templates for both files from the *uORF-Tools/templates* into the *uORF-Tools/* folder.

```
$ cp uORF-Tools/templates/fastq-samples.tsv uORF-Tools/
```

The template looks as follows:

| **method** | **condition** | **replicate** | **inputFile** |
|---|---|---|---|
| RNA | A | 1 | fastq/SRR6026769.fastq.gz |
| RNA | A | 2 | fastq/SRR6026770.fastq.gz |
| RNA | B | 1 | fastq/SRR6026765.fastq.gz |
| RNA | B | 2 | fastq/SRR6026766.fastq.gz |
| RIBO | A | 1 | fastq/SRR6026777.fastq.gz |
| RIBO | A | 2 | fastq/SRR6026778.fastq.gz |
| RIBO | B | 1 | fastq/SRR6026773.fastq.gz |
| RIBO | B | 2 | fastq/SRR6026774.fastq.gz |

**Please ensure not to replace any tabulator symbols with spaces while changing this file.**

Next, we are going to set up the *config.yaml*.

```
$ cp uORF-Tools/templates/config.yaml uORF-Tools/
$ vi uORF-Tools/config.yaml
```

This file contains the following variables:

- **taxonomy** Specify the taxonomic group of the used organism in order to ensure the correct removal of reads mapping to ribosomal genes (Eukarya, Bacteria, Archea). (Option for the preprocessing workflow)
- **adapter** Specify the adapter sequence to be used. If not set, *Trim galore* will try to determine it automatically. (Option for the preprocessing workflow)
- **samples** The location of the samples sheet created in the previous step.
- **genomeindexpath** If the STAR genome index was already precomputed, you can specify the path to the files here, in order to avoid recomputation. (Option for the preprocessing workflow)
- **uorfannotationpath** If a uORF-annotation file was already pre-computed, you can specify the path to the file here. Please make sure, that the file has the same format as the uORF_annotation_hg38.csv file provided in the git repo (i.e. same number of columns, same column names)
- **alternativestartcodons** Specify a comma separated list of alternative start codons.

Change the config file as shown in Figure 6. For this tutorial, we can keep the

```
#Taxonomy of the samples to be processed, possible are
    Eukarya, Bacteria, Archea
taxonomy: "Eukarya"
#Adapter sequence used
adapter: ""
samples: "uORF-Tools/fastq-samples.tsv"
genomeindexpath: ""
uorfannotationpath: ""
alternativestartcodons: "CTG,GTG,TTG"
```

**Fig. 6.** The contents of the configuration file (*config.yaml*)

default values for the *config.yaml* as shown in Figure 6. The organism analyzed in this tutorial is *homo sapiens*, therefore we keep the taxonomy at *Eukarya*. The path to *samples.tsv* is set correctly.

#### 8 Running the workflow

Now that we have all the required files, we can start running the workflow, either locally or in a cluster environment.

**Information about processing parameters** In this pipeline we use trim_galore for adapter and quality trimming (Parameters: –phred33 -q 20 –length 15 –trim-n –suppress_warn –clip_R1 1), sortmerna for rRNA removal (rRNA fasta files are obtained from
https://github.com/biocore/sortmerna/tree/master/rRNA_databases
, according to the specified Taxon (i.e. Eukarya, Bacteria, Archea)) and STAR for read alignment.
(Parameters: –outSAMtype BAM SortedByCoordinate
–outSAMattributes All –outFilterMultimapNmax 1
–alignEndsType Extend5pOfRead1).

**Run the workflow locally** Use the following steps when you plan to execute the workflow on a single server or workstation. Please be aware that some steps of the workflow require a lot of memory, specifically for eukaryotic species.

```
$ snakemake --use-conda -s uORF-Tools/Preprocessing_Snakefile\
            --configfile uORF-Tools/config.yaml \
            --directory ${PWD} -j 20 \
            --latency-wait 60
```

**Run Snakemake in a cluster environment** Use the following steps if you are executing the workflow via a queuing system. Edit the configuration file cluster.yaml according to your queuing system setup and cluster hardware. The following system call shows the usage with Grid Engine:

```
$ snakemake --use-conda -s uORF-Tools/Preprocessing_Snakefile\
       --configfile uORF-Tools/config.yaml \
       --directory ${PWD} -j 20 \
       --cluster-config uORF-Tools/templates/sge-cluster.yaml
```

```
#!/bin/bash
#PBS -N <ProjectName>
#PBS -S /bin/bash
#PBS -q "long"
#PBS -d <PATH/ProjectFolder>
#PBS -l nodes=1:ppn=1
#PBS -o <PATH/ProjectFolder>
#PBS -j oe
cd <PATH/ProjectFolder>
source activate snakemake
snakemake --latency-wait 600 --use-conda -s uORF-Tools/
    Snakefile --configfile uORF-Tools/config.yaml --
    directory ${PWD} -j 20 --cluster-config uORF-Tools/
    torque.yaml --cluster "qsub -N {cluster.jobname} -S /
    bin/bash -q {cluster.qname} -d <PATH/ProjectFolder> -l
     {cluster.resources} -o {cluster.logoutputdir} -j oe"
```

**Fig. 7.** Bash script to run the example workflow in a TORQUE cluster environment.

**Example: Run Snakemake in a cluster environment** We ran the tutorial workflow in a cluster environment, specifically a TORQUE cluster environment. Therefore, we created a bash script *torque.sh* in our project folder.

```
$ vi torque.sh
```

We proceeded by writing the queueing script as shown in Figure 7.
We then simply submitted this job to the cluster.

```
$ qsub torque.sh
```

Using any of the presented methods, this will run the workflow on our dataset and create the desired output files.

### Bibliography

[1] Johannes Köster and Sven Rahmann. SnakemakeâĂŤa scalable bioinformatics workflow engine. *Bioinformatics*, page bty350, 2018.

[2] Björn Grüning, Ryan Dale, Andreas Sjödin, Jillian Rowe, Brad A. Chapman, Christopher H. Tomkins-Tinch, Renan Valieris, and Johannes Köster. Bioconda: A sustainable and comprehensive software distribution for the life sciences. *bioRxiv*, 2017.

[3] J. Harrow, A. Frankish, J. M. Gonzalez, E. Tapanari, M. Diekhans, F. Kokocinski, B. L. Aken, D. Barrell, A. Zadissa, S. Searle, I. Barnes, A. Bignell, V. Boychenko, T. Hunt, M. Kay, G. Mukherjee, J. Rajan, G. Despacio-Reyes, G. Saunders, C. Steward, R. Harte, M. Lin, C. Howald, A. Tanzer, T. Derrien, J. Chrast, N. Walters, S. Balasubramanian, B. Pei, M. Tress, J. M. Rodriguez, I. Ezkurdia, J. van Baren, M. Brent, D. Haussler, M. Kellis, A. Valencia, A. Reymond, M. Gerstein, R. Guigo, and T. J. Hubbard. GENCODE: the reference human genome annotation for The ENCODE Project. *Genome Res.*, 22(9):1760–1774, Sep 2012.

[4] Daniel R Zerbino, Premanand Achuthan, Wasiu Akanni, MÂăRidwan Amode, Daniel Barrell, Jyothish Bhai, Konstantinos Billis, Carla Cummins, Astrid Gall, Carlos GarcÃŋa GirÃşn, Laurent Gil, Leo Gordon, Leanne Haggerty, Erin Haskell, Thibaut Hourlier, Osagie G Izuogu, Sophie H Janacek, Thomas Juettemann, Jimmy Kiang To, Matthew R Laird, Ilias Lavidas, Zhicheng Liu, Jane E Loveland, Thomas Maurel, William McLaren, Benjamin Moore, Jonathan Mudge, Daniel N Murphy, Victoria Newman, Michael Nuhn, Denye Ogeh, Chuang Kee Ong, Anne Parker, Mateus Patricio, Harpreet Singh Riat, Helen Schuilenburg, Dan Sheppard, Helen Sparrow, Kieron Taylor, Anja Thormann, Alessandro Vullo, Brandon Walts, Amonida Zadissa, Adam Frankish, Sarah E Hunt, Myrto Kostadima, Nicholas Langridge, Fergal J Martin, Matthieu Muffato, Emily Perry, Magali Ruffier, Dan M Staines, Stephen J Trevanion, Bronwen L Aken, Fiona Cunningham, Andrew Yates, and Paul Flicek. Ensembl 2018. *Nucleic Acids Research*, 46(D1):D754–D761, 2018.
