## Supplementary Table S1 for "*uORF-Tools* – Workflow for the determination of translation-regulatory upstream open reading frames"

**Table S1.** Ribo-seq data series used for the generation of the comprehensive human uORF annotation file. Data sets came from nine different data series and consisted of variable numbers of treatments and replicates. 35 data sets of individual samples each included ribo-seq and associated RNA-seq data.

| Data series # | Date series ID | Data set (ribo-seq IDs) |
|---|---|---|
| 1 | GSE103719 | GSM2779669 / GSM2779661<br>GSM2779670 / GSM2779662<br>GSM2779671 / GSM2779663<br>GSM2779672 / GSM2779664<br>GSM2779673 / GSM2779665<br>GSM2779674 / GSM2779666<br>GSM2779675 / GSM2779667<br>GSM2779676 / GSM2779668 |
| 2 | GSE66929 | GSM1634443 / GSM1632189<br>GSM1634445 / GSM1632191<br>GSM1634449 / GSM1632193 |
| | GSE96716 | GSM2538903 / GSM2538901<br>GSM2538904 / GSM2538902 |
| 4 | GSE42509 | GSM1047584 / GSM1041191<br>GSM1047585 / GSM1041192<br>GSM1047586 / GSM1041193<br>GSM1047587 / GSM1041194<br>GSM1047591 / GSM1041199 |
| 5 | GSE69602 | GSM1704511 / GSM1704559<br>GSM1704513 / GSM1704561<br>GSM1704523 / GSM1704571<br>GSM1704525 / GSM1704573 |
| 6 | GSE56924 | GSM1371443 / GSM1371395<br>GSM1371449 / GSM1371401<br>GSM1371455 / GSM1371407<br>GSM1371461 / GSM1371413 |
| 7 | GSE96714 | GSM2538884 / GSM2538879 |
| 8 | GSE114636 | GSM3146275 / GSM3146283<br>GSM3146276 / GSM3146284 |
| 9 | GSE98623 | GSM2602082 / GSM2602073<br>GSM2602083 / GSM2602074<br>GSM2602084 / GSM2602075<br>GSM2602085 /GSM2602076<br>GSM2602086 / GSM2602077<br>GSM2602087 / GSM2602078 |
